# Deletion of Rap-Phr systems in *Bacillus subtilis* influences *in vitro* biofilm formation and plant root colonization

**DOI:** 10.1101/2021.03.15.435437

**Authors:** Mathilde Nordgaard Christensen, Rasmus Møller Rosenbek Mortensen, Nikolaj Kaae Kirk, Ramses Gallegos-Monterrosa, Ákos T. Kovács

**Affiliations:** Bacterial Interactions and Evolution Group, DTU Bioengineering, Technical University of Denmark, 2800 Kongens Lyngby, Denmark; Terrestrial Biofilms Group, Institute of Microbiology, Friedrich-Schiller-University Jena, 07743 Jena, Germany

**Keywords:** *Bacillus subtilis*, Rap-Phr, matrix gene expression, biofilm formation, root colonization

## Abstract

Natural isolates of the soil-dwelling bacterium *Bacillus subtilis* form robust biofilms under laboratory conditions and colonize plant roots. *B. subtilis* biofilm gene expression displays phenotypic heterogeneity that is influenced by a family of Rap-Phr regulatory systems. Most Rap-Phr systems in *B. subtilis* have been studied independently, in different genetic backgrounds and under distinct conditions, hampering true comparison of the Rap-Phr systems’ impact on bacterial differentiation. Here, we investigated each of the 12 Rap-Phr systems of *B. subtilis* NCIB 3610 for their effect on biofilm formation. By studying single Δ*rap-phr* mutants, we show that despite redundancy between the cell-cell communication systems, deletion of each of the 12 Rap-Phr systems influences matrix gene expression, which could possibly enable fine-tuning of the timing and level of matrix production in response to specific conditions. Furthermore, some of the Δ*rap-phr* mutants demonstrated altered biofilm formation *in vitro* and colonization of *Arabidopsis thaliana* roots, but not necessarily similarly in both processes, indicating that the pathways regulating matrix gene expression and other factors important for biofilm formation may be differently regulated under these distinct conditions.

**Significance Statement:** Natural isolates of *Bacillus subtilis* form robust biofilms *in vitro* and on plant roots. The formation of these heterogeneous populations is regulated by diverse Rap-Phr systems. However, most Rap-Phr systems in *B. subtilis* have been studied independently and in different genetic backgrounds. Here, we report that all 12 Rap-Phr systems affect matrix gene expression, while some of them affect development of *in vitro* biofilms and plant root colonization. Our study highlights the importance of the Rap-Phr systems in environmental adaptation of *B. subtilis*, specifically during biofilm formation in the rhizosphere.

## Introduction

In nature, biofilms are the predominant lifestyle of bacteria and are known as surface-associated microbial communities embedded in a self-produced matrix (Hall-Stoodley *et al*., 2004; Flemming and Wingender, 2010; López *et al*., 2010). Biofilms have been widely studied since they represent a fascinating example of microbial development in response to environmental cues (O’Toole *et al*., 2000). Furthermore, studying biofilms is of special interest due to their detrimental impact in clinical and industrial settings (Stewart, 2002; Di Pippo *et al*., 2018) as well as their promising potential within the biotechnology industry (Singh *et al*., 2006; Blake *et al*., 2021). Regarding the latter, the Gram-positive bacterium *Bacillus subtilis* has in the last two decades gained interest due to its promising potential as a biocontrol agent within agriculture (Ongena and Jacques, 2008; Kiesewalter *et al*., 2021). In its natural habitat, the soil-dwelling bacterium colonizes plants by forming a biofilm on the root (Bais *et al*., 2004; Chen *et al*., 2012; Beauregard *et al*., 2013). After successfully colonizing the root, *B. subtilis* exerts its plant-beneficial properties, including directly promoting plant growth as well as protecting the plant against diseases (Blake *et al*., 2021). Furthermore, *B. subtilis* forms spores that are highly resistant to extreme environments (Piggot and Hilbert, 2004) facilitating easy formulation and storage of *B. subtilis* for application within agriculture (Ongena and Jacques, 2008).

*B. subtilis* can easily be isolated from the rhizosphere of plants (Fall *et al*., 2004), and a study performed by Chen *et al*. (2013) showed that the majority of natural strains isolated from rhizosphere formed architecturally complex biofilms under laboratory conditions, indicating that biofilm formation is an important trait for *B. subtilis* to thrive in its natural habitat. In the laboratory, *B. subtilis* has long been studied using different kinds of biofilm models including colonies at the air-agar interface as well as floating biofilms formed at the air-liquid interface, termed pellicles (Arnaouteli *et al*., 2021). A prevalent feature of *B. subtilis* biofilms is that they display complex phenotypic heterogeneity, where genetically identical cells differentiate into distinct cell types in response to external cues (Lopez *et al*., 2009; López and Kolter, 2010). The variation in environmental conditions throughout the biofilm (Costerton *et al*., 1994) thereby leads to a heterogeneous population with different cell types performing distinct tasks and occupying different micro-niches. The extracellular signals triggering cell differentiation include quorum-sensing molecules, natural products, and nutrient availability that activate a set of sensor kinases (Mhatre *et al*., 2014; Arnaouteli *et al*., 2021). Once activated, the sensor kinases phosphorylate their respective master transcriptional regulators, Spo0A, DegU and ComA, each of which activates different sets of genes (López and Kolter, 2010). The Spo0A pathway governs differentiation into matrix-producing cells as well as sporulating cells. In response to external cues, one or more of five histidine kinases, KinA-E, are activated which results in phosphorylation of Spo0F. Spo0F∼P then transfers its phosphoryl group to Spo0B, which in turn transfers the phosphoryl group to and thereby activates Spo0A (Jiang *et al*., 2000b; Fujita *et al*., 2005). At low Spo0A∼P levels, the genes involved in the synthesis of matrix components, exopolysaccharide (EPS) and TasA protein fiber, are expressed (Fujita *et al*., 2005; Cairns *et al*., 2014). These two matrix components are well known to be required for biofilm formation *in vitro* and on the plant root (Branda *et al*., 2006; Beauregard *et al*., 2013). When high levels of Spo0A∼P are reached, genes involves in sporulation are expressed (Fujita *et al*., 2005). The DegU response regulator is phosphorylated by its cognate histidine kinase DegS. Studies have indicated that inhibition of flagellar rotation, as may take place upon contact with a surface, acts as a mechanical trigger to activate the DegS-DegU two-component signaling pathway (Cairns *et al*., 2013). At very low levels of DegU∼P, genes related to swarming motility are expressed, while elevated levels of DegU∼P induce exoprotease production and at the same time represses motility genes (Verhamme *et al*., 2007; Belas, 2013). Finally, the pheromone ComX activates the histidine kinase ComP, which phosphorylates ComA, resulting in expression of genes involved in competence development and surfactin production (Comella and Grossman, 2005).

This regulatory network governing cell differentiation in *B. subtilis* is further regulated by a family of response regulator aspartyl-phosphate (Rap) phosphatases and their associated phosphatase regulator (Phr) peptides (Perego, 2013). In the *B. subtilis* group, 80 distinct putative *rap-phr* alleles have been identified with a strain having on average 11 *rap* genes (Even-Tov *et al*., 2016). The abundance of Rap and Phr peptides is transcriptionally controlled in response to different cellular signals (Mueller *et al*., 1992; Perego *et al*., 1994; Lazazzera *et al*., 1999; Jiang *et al*., 2000a; Jarmer *et al*., 2001; McQuade *et al*., 2001; Ogura *et al*., 2001; Auchtung *et al*., 2005). The genes encoding the Rap-Phr pairs are found as gene cassettes with the *phr* gene immediately downstream of the *rap* gene and the expression of these being transcriptionally coupled, with some *phr* genes also being transcribed independently of their cognate *rap* genes from promoters controlled by σ^H^ (Reizer *et al*., 1997; Pottahil and Lazazzera, 2003; McQuade *et al*., 2001). Some exceptions to this exist, for example the *rapB* gene is not followed by an active peptide encoding gene (Perego *et al*., 1996). Moreover some Rap proteins are regulated by Phr peptides encoded in other cassettes, e.g. RapB and J are both controlled by PhrC (Parashar, Jeffrey, *et al*., 2013). When expressed, the Rap phosphatases exert their effect within the cell by either dephosphorylating Spo0F∼P (thus hindering Spo0A phosphorylation) or inhibiting the DNA binding activity of ComA or DegU (Perego, 2013). In contrast, the product of the *phr* gene is secreted out of the cell through the Sec-dependent export pathway and processed into mature five to six amino acid signaling peptides. At high cell density, the Phr peptides reach threshold concentrations at which they are transported back into the cell by the oligopeptide permease (Opp) (Pottahil and Lazazzera, 2003; Perego 2013). Once within the cell, the Phr peptides will inhibit their cognate Rap proteins, thereby relieving the inhibition of the master regulators resulting in altered gene expression (Pottahil and Lazazzera, 2003; Perego, 2013). The Rap-Phr systems thereby act as cell-cell signaling systems in *B. subtilis*, allowing the bacteria to respond to environmental changes only at sufficient cell densities.

As expected by the diversity and abundance of multiple Rap-Phr systems regulating the activity of these three master regulators, the Rap phosphatases show high redundancy in their regulatory function: RapA, B, E, H, I, J and P have been shown to dephosphorylate Spo0F∼P, RapC, D, F, H, K and P regulate ComA, while RapG has been shown to regulate the activity of DegU (Auchtung *et al*., 2006; Ogura and Fujita, 2007; Perego, 2013; Omer Bendori *et al*., 2015) (Table S1). Furthermore, RapI is involved in the regulation of mobile genetic elements, as it activates the propagation of the mobile genetic element that encodes it (Auchtung *et al*., 2005). The regulation of the master regulators by multiple Rap phosphatases allows the integration of diverse signals to control cell differentiation in response to different conditions. However, this overall overview of the Rap-Phr signaling network in *B. subtilis* is based on studies where most Rap-Phr systems have been tested in different genetic backgrounds and under distinct cultivation conditions (Perego, 2013). Additionally, previous investigations in *B. subtilis* have directed their study towards certain targets of Rap-Phr regulation, with RapA and B being mostly studied for their impact on sporulation (Perego and Hoch, 1996), while RapC and F have been shown to be involved in competence development (Core and Perego, 2003; Bongiorni *et al*., 2005). So far, only RapP has been demonstrated to impact biofilm formation (Parashar, Konkol, *et al*., 2013; Omer Bendori *et al*., 2015). We previously studied all 12 Rap-Phr systems of the undomesticated strain *B. subtilis* NCIB 3610 in the same genetic background by following the relative abundance of all possible single and double Δ*rap-phr* mutants as well as the WT (79 strains) in populations subjected to different selective conditions. This study highlighted that the variability in Rap-Phr systems affected the ability to compete in diverse environments (Gallegos-Monterrosa *et al*., 2021).

In this study, we systematically investigated the contribution of each of the 12 Rap-Phr systems in *B. subtilis* 3610 to biofilm formation. We assessed wild type (WT) and the 12 single Δ*rap-phr* mutants for matrix gene expression and biofilm formation under different conditions. We found that all 12 mutants showed altered matrix gene expression compared to the WT. Furthermore, we observed that the Rap-Phr modules not only affect *in vitro* biofilm formation but also colonization of plant root that represents an ecologically relevant environment.

## Results

*Single Δ*rap-phr *mutants show altered matrix gene expression compared to the WT* To study the effect of Rap-Phr systems of *B. subtilis* on biofilm formation, we used the *B. subtilis* DK1042 strain (a natural competent derivative of the undomesticated NCBI 3610) (Konkol *et al*., 2013) as WT. In contrast to domesticated strains which may have acquired mutations to the Rap-Phr systems or even lost some of them, NCIB 3610 and the derived DK1042 (hereafter referred to as 3610) contain all 12 *rap-phr* modules (McLoon *et al*., 2011). Rap-Phr systems of *B. subtilis* have been studied using both overexpression as well as deletion mutants of the Rap-Phr systems as reviewed by Perego *et al*. (2013). Interestingly, the absence of Rap modules in a natural isolate of *B. subtilis* was shown to influence sporulation timing (Serra *et al*., 2014). In this study, we employed single Δ*rap-phr* mutants of *B. subtilis* created in our previous work (Gallegos-Monterrosa *et al*., 2021) to study the impact of the Rap-Phr modules on biofilm formation. The diversity in Rap-Phr regulatory systems as well as the reported role of several of the Rap phosphatases in regulating the activity of Spo0A (Perego, 2013; Omer Bendori *et al*., 2015), which controls matrix gene expression, prompted us to test WT and the 12 single Δ*rap-phr* mutants for matrix gene expression. To quantify matrix gene expression during biofilm formation, we used a previously established approach (Kearns *et al*., 2005). This method utilizes the observation that expression from the promoter of the *tapA-sipW-tasA* operon, responsible for the production of the protein component of the extracellular matrix (Branda *et al*., 2006), is highly induced during late exponential growth phase under shaking conditions in MSgg, a medium known to induce biofilm formation. After 6h of growth in MSgg under shaking conditions, the fluorescence intensities of WT and mutants harboring a P_*tapA*_-*gfp* reporter construct were measured using flow cytometry. Importantly, the side scatter (SSC-A) vs forward scatter (FSC-A) plots revealed no big changes in cell granularity or size of the mutants compared to the WT (Fig. S1). Since RapA, B, E, H, I, J and P have been reported to dephosphorylate Spo0F∼P (Perego, 2013; Omer Bendori *et al*., 2015), we hypothesized that the mutants lacking one of these Rap phosphatases would have a larger relative ON population (i.e. cells in the high expression state), due to more cells committing to matrix production, and/or display a higher mean expression from P_*tapA*_ in the ON population due to earlier matrix gene expression, as compared to the WT. Alternatively, if redundancy operates between some of the Rap-Phr systems, the absence of one Rap-Phr system may have only minor or insignificant effects due to the expression and function of another redundant system. However, *ΔrapA* showed a significantly smaller relative ON population (p=0.022, n=3-8) than the WT (Fig. 1), while all other mutants showed a significantly higher mean expression of the ON population compared to the WT. The increased mean expression by the mutants could potentially be caused by enhanced growth in MSgg, as strains growing faster would reach the threshold density for biofilm formation earlier, and thus show increased matrix gene expression. However, when the optical density (OD) at 600 nm was measured after 6 h in MSgg prior to flow cytometry analysis, the mutants showed either similar or reduced OD_600_ compared to WT (data not shown), thus excluding this explanation. Interestingly, the increased matrix gene expression was not limited to the mutants lacking one of the Rap phosphatases reported to dephosphorylate Spo0F∼P activity. These results show that all 12 Rap-Phr regulatory systems influence matrix gene expression under these conditions.

**Fig. 1.**
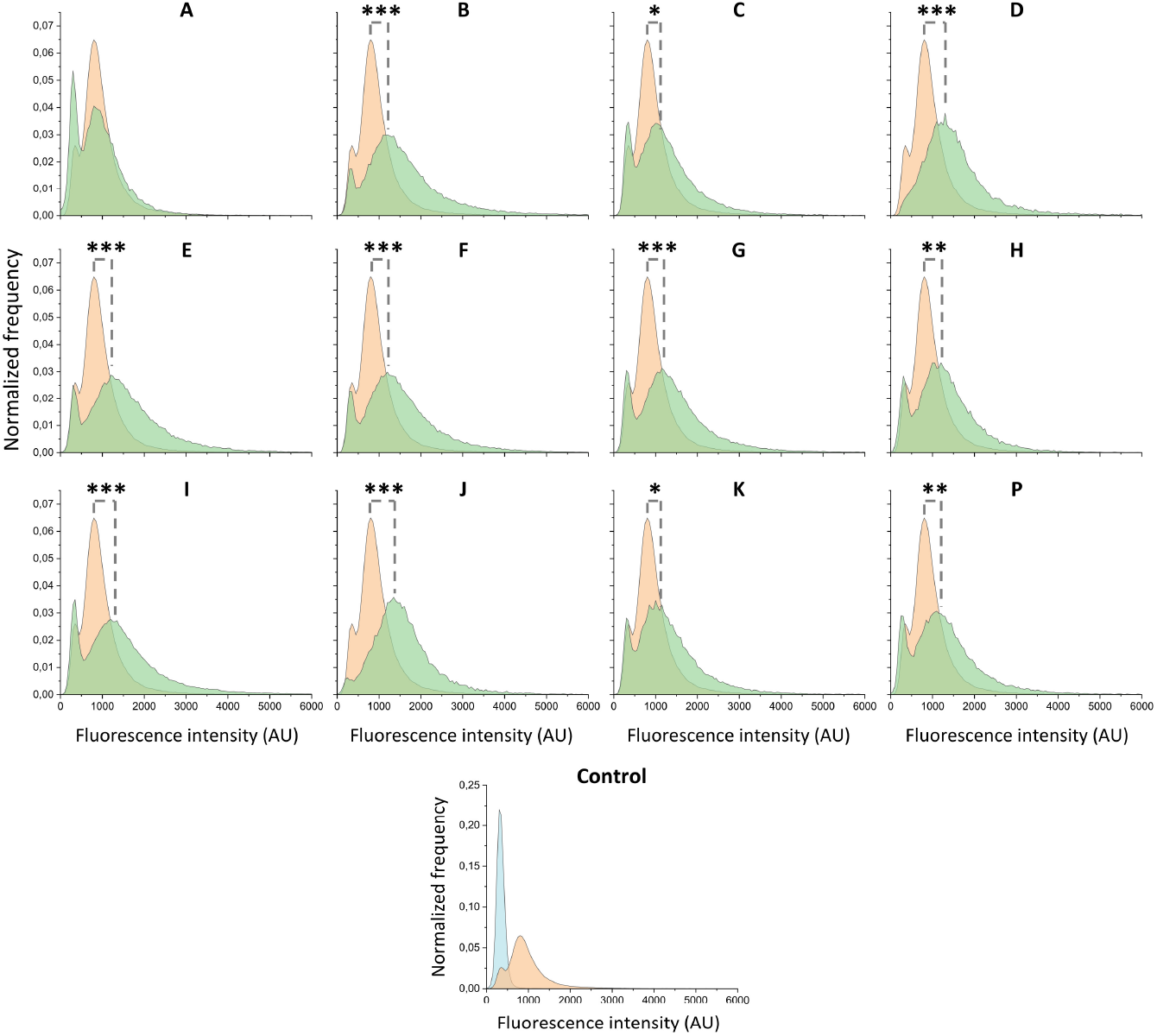
Expression of the matrix gene *tapA* in *B. subtilis* WT and Δ*rap-phr* mutants after growth in MSgg under shaking conditions. Flow cytometry analysis showing the average distributions of GFP expression of Δ*rap-phr* mutants and WT harboring the P_*tapA*_*-gfp* construct (n=3-8). The average WT distribution is shown in each graph for comparison. Orange = WT, green = mutant, blue = non-labelled control strain. Letters denote the corresponding Δ*rap-phr* mutants, i.e. A = Δ*rapA*, B = Δ*rapB* and so forth. AU indicates arbitrary units. Significant difference in the mean fluorescence intensity of the ON population (GFP values between 500 and 10.000) between mutants and WT were tested by an ANOVA followed by Dunnett’s Multiple Comparison test. *: P<0.05, **: P<0.01, ***: P<0.001.

### Absence of certain Rap-Phr modules alters biofilm formation in vitro

Since matrix production is required for proper biofilm development (Branda *et al*., 2006; Dragoš *et al*., 2018), and most Δ*rap-phr* mutants showed increased matrix gene expression under shaking conditions in MSgg, we hypothesized that this would manifest in more complex and robust biofilm formation. To investigate this, WT and mutants were tested for their ability to form biofilm on a solid surface (MSgg agar) and robust pellicle biofilms at the MSgg medium-air interface. The pellicle forms as oxygen in the medium is exhausted and *B. subtilis* moves towards higher oxygen concentrations, thus the air-liquid surface, where cells create a biofilm (Hölscher *et al*., 2015).

In accordance to previous work, *B. subtilis* WT produced a wrinkled colony biofilm, as well as a robust, wrinkled pellicle (Branda *et al*., 2001; Gallegos-Monterrosa *et al*., 2016) (Fig. 2). While Δ*rapA, B, J* and *K* formed more wrinkled colonies, Δ*rapI* and *P* formed complex but very small colonies on MSgg agar compared to the WT. In contrast, Δ*rapC* formed a large, transparent colony with fewer wrinkles than WT. When testing for development of pellicle biofilms, the Δ*rapA, C, I* and *P* mutants formed thin and/or non-homogenous pellicles, while the qualitative assessment revealed no mutants with more complex or robust pellicles than WT (Fig. 2). Lastly, the Δ*rapD, E, F, G* and *H* mutants formed comparable colony and pellicle biofilms to the WT. These results show that despite most *Δrap-phr* mutants showed increased matrix gene expression under shaking conditions compared to the WT, this did not necessarily manifest in development of more complex biofilms on agar or at the air-liquid interface.

**Fig. 2.**
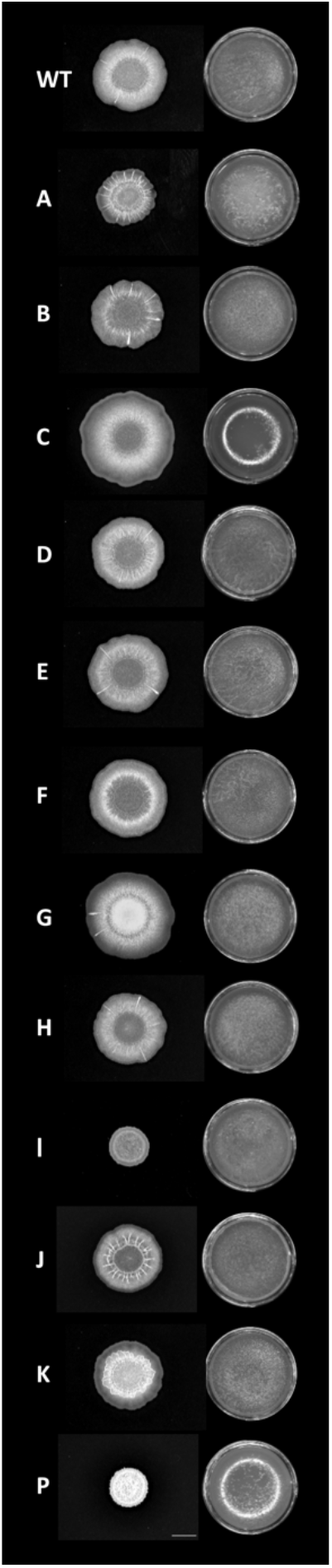
Biofilm formation of *B. subtilis* WT and single Δ*rap-phr* mutants. Overnight cultures of *B. subtilis* WT and Δ*rap-phr* mutants were spotted on MSgg medium solidified with 1.5 % agar (left) or inoculated in MSgg with a starting OD_600_ = 0.05 (right). Images were taken from above after 48 h of incubation at 30°C using a stereomicroscope. Letters denote the corresponding Δ*rap* mutant, i.e. A = Δ*rapA*, B = Δ*rapB* and so forth. Bar denotes 5 mm for the biofilm colony images.

### Certain Δrap-phr mutants are affected in colonization of Arabidopsis thaliana roots

To reveal how the Rap-Phr modules affect biofilm formation in a more ecologically relevant environment, WT and mutants were tested for biofilm formation on the roots of the model plant organism *A. thaliana*. Similar to biofilm formation *in vitro* (Branda *et al*., 2006), previous studies have shown that matrix production is required for biofilm formation on *A. thaliana* roots (Beauregard *et al*., 2013; Dragoš *et al*., 2018). We therefore speculated that the increased matrix gene expression observed in most strains would enable enhanced biofilm formation on plant roots.

For this purpose, sterile *A. thaliana* seedlings with a root length of 0.5 to 1.2 cm were inoculated with one of the 12 Δ*rap-phr* mutants or WT and incubated in a plant chamber at 24°C (90 rpm) for 16 h, after which root colonization was quantified as colony forming unit (CFU) per mm root length. Of the 12 mutants tested, only Δ*rapD, J* and *P* mutants showed significantly increased root colonization compared to the WT (Fig. 3). In contrast, the *ΔrapI* mutant was significantly reduced in root colonization. The observation that Δ*rapI* and *P* showed similar biofilm formation *in vitro* (Fig. 2) but performed oppositely during root colonization (Fig. 3) might indicate that the effect of each individual Rap phosphatase on biofilm formation depends on the environment. Furthermore, similarly to biofilm formation *in vitro*, these results show that increased matrix gene expression under shaking conditions did not necessarily manifest in increased root colonization.

**Fig. 3.**
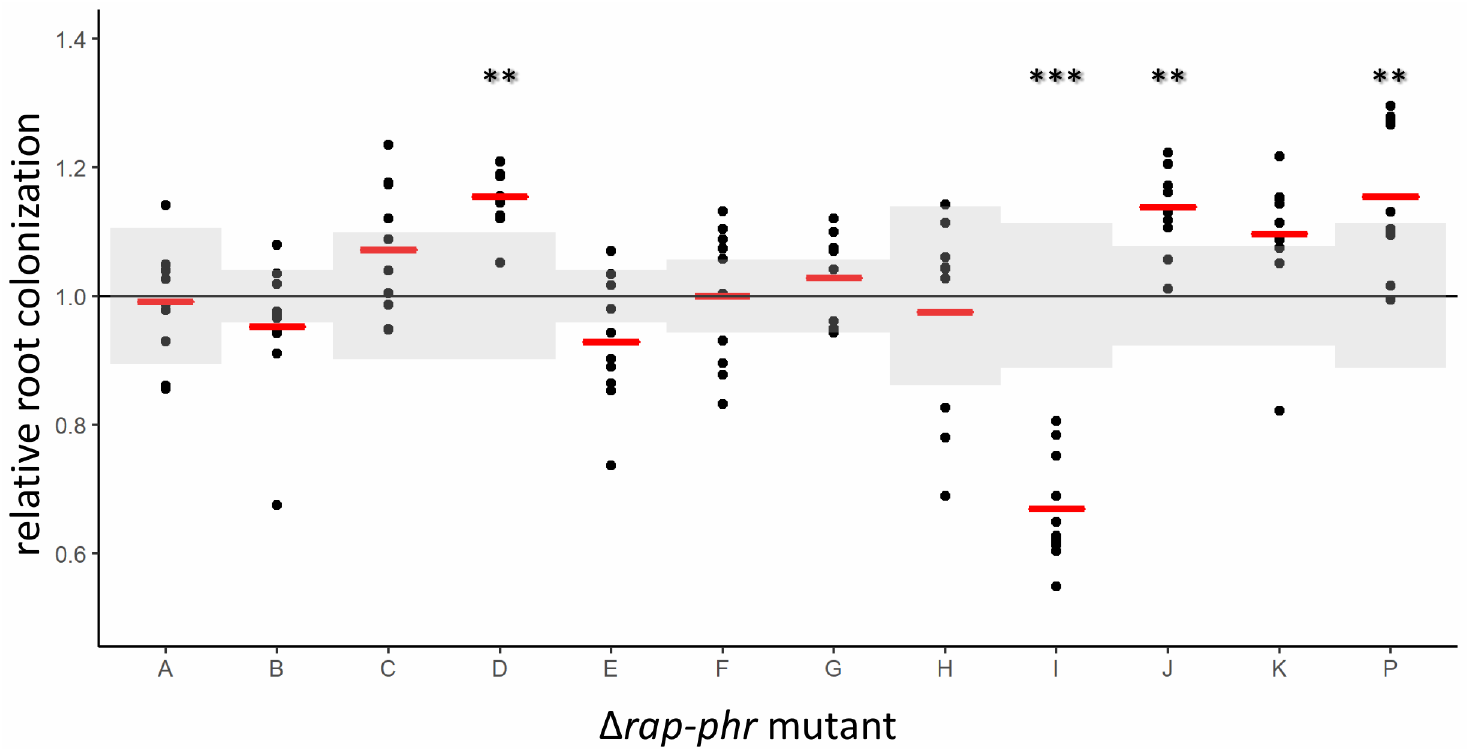
*A. thaliana* root colonization by *B. subtilis* WT and single Δ*rap-phr* mutants. To estimate the impact of each of the Rap-Phr modules on root colonization, WT and single Δ*rap-phr* mutants were inoculated onto five-day-old seedlings of *A. thaliana* (n=7-10). After 16 h, CFU per mm root length was quantified. The log10-transformed value of CFU/mm root for each technical replicate was normalized to the mean of the WT from the same day. Each dot represents a root, while the mean value for each mutant is displayed as a red horizontal line. The black horizontal line represents the mean of the WT and the SD of the WT from each respective experiment is shown in shaded grey. For each assay, an ANOVA was performed on the log10-transformed values of CFU/mm root length. When significant (P<0.05), means were compared via Dunnett’s multiple comparison test with WT as the control. When data failed to meet parametric assumptions, a Kruskal-Wallis test was performed followed by a Dunn’s test. *: P<0.05, **: P<0.005, ***: P<0.001.

### Certain Δrap-phr mutants show altered growth compared to WT

The timing of biofilm initiation *in vitro* and on the root depends on the cell density, as only at sufficiently high cell density the Phr peptides will reach threshold concentrations allowing them to be imported into the cell, where they will inhibit their cognate Rap phosphatases. Consequently, a subset of the Rap phosphatases influences phosphorylation of Spo0A and expression of biofilm genes. Biofilm formation is therefore affected by growth rate, as strains growing faster will reach the threshold density for biofilm formation earlier. In order to test whether the observed changes in biofilm formation *in vitro* and on the root could be (partly) attributed to altered growth, the 12 Δ*rap-phr* mutants were tested for growth in MSgg under shaking conditions. For clear visualization, the growth curves of the 12 mutants were separated into four plots (Fig. 4). Of the 12 mutants tested, only Δ*rapA, I* and *P* showed altered growth profile compared to WT. Δ*rapA* and *I* showed a reduced growth rate, a delayed entry into stationary phase and reduced max OD_590_. In addition, Δ*rapI* had a prolonged stationary phase as well as slower decline during late stationary phase (possibly death phase). Δ*rapP* showed slower growth during exponential phase, which continued for about 10 h longer than the WT, but displayed a higher max OD_590_ at stationary phase. In LB medium, the same mutants were similarly or less affected in growth compared to WT (Fig. S2). The impaired growth of Δ*rapA* and *I* is in accordance with these two mutants forming thin and/or non-homogeneous pellicles (Fig.2), and Δ*rapI* displaying reduced root colonization (Fig. 3). In contrast, the prolonged exponential growth of Δ*rapP* is inconsistent with the thin pellicle formed by this mutant, but correlating with the increased root colonization observed for this mutant. Finally, Δ*rapC* showed similar growth profile as the WT but formed a thin pellicle. The mutants have differential biofilm forming capacity *in vitro* and, on the root, which, interestingly, cannot be directly correlated with the growth rates and profiles of the mutants.

**Fig. 4.**
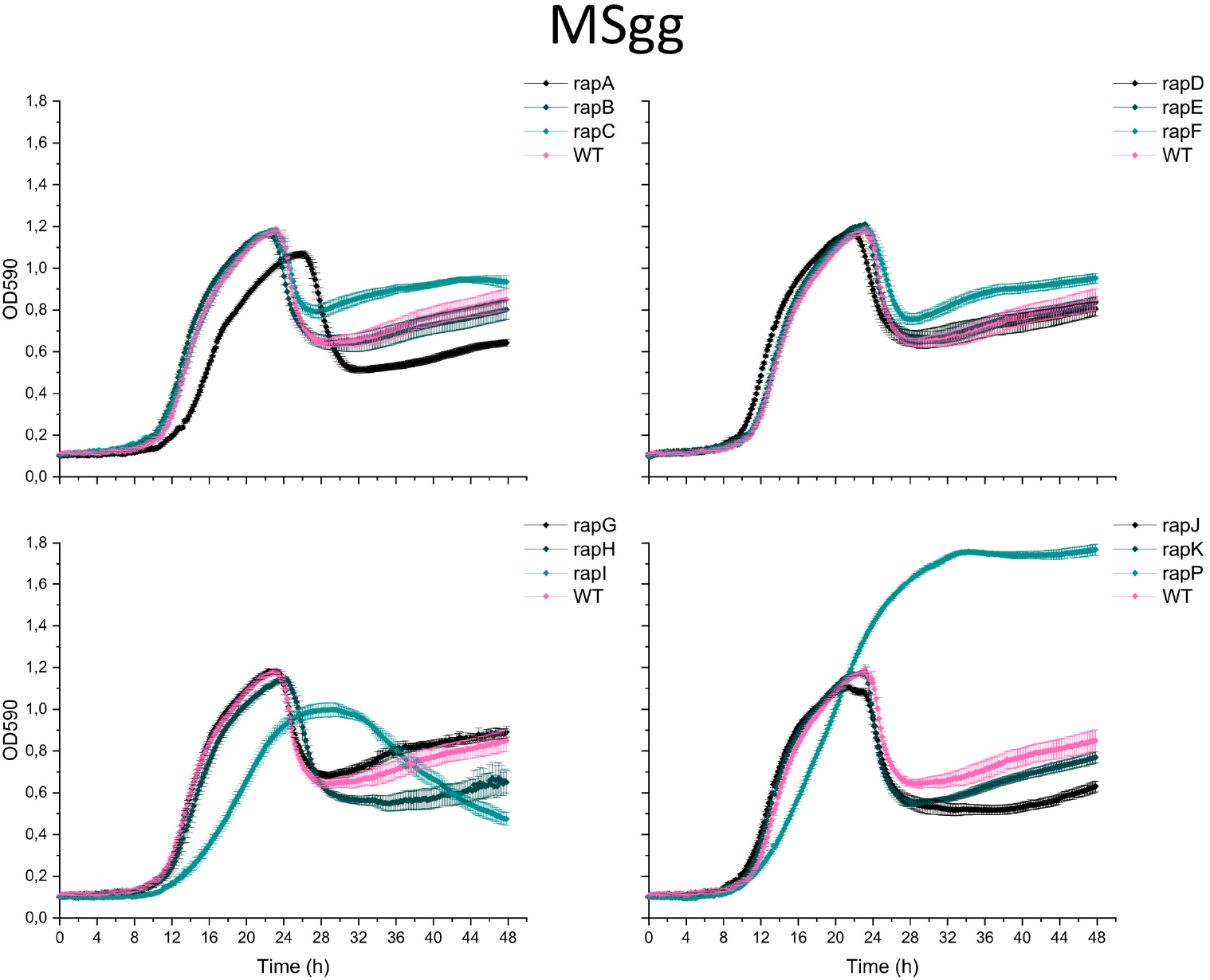
Growth of *B. subtilis* WT and single Δ*rap-phr* mutants in MSgg. WT and mutants were inoculated into 96-well plates with a starting OD_590_ of 0.05. OD_590_ was measured every 10 min for 48 h at 30°C; each time point represents the mean of six technical replicates from two overnight cultures (N=6). Error bars represent standard error (SE).

## Discussion

For decades, the Rap-Phr regulatory systems which control the activity of the three master regulators governing cell differentiation in *B. subtilis* have been extensively studied (Pottahil and Lazazzera, 2003; Perego, 2013). However, since most studies have investigated the Rap-Phr networks of *B. subtilis* independently from each other, in different genetic backgrounds and under distinct conditions, it is difficult to compare the results from these studies. Here, we investigated all 12 Rap-Phr systems found in *B. subtilis* 3610 for their effect on matrix gene expression and biofilm development *in vitro*. In addition, we examined the impact of Rap-Phr systems in colonization of plant roots for the first time.

Several of the Rap phosphatases have been reported to dephosphorylate Spo0F∼P (Perego, 2013; Omer Bendori *et al*., 2015) which is expected to influence matrix production, but only RapP has previously been demonstrated to affect matrix gene expression (Parashar, Konkol, *et al*., 2013; Omer Bendori *et al*., 2015). We were therefore interested in testing the effect of each of the 12 Rap-Phr systems on matrix gene expression. Inspired by a previously established method (Kearns *et al*., 2005), expression from the promoter of the *tapA-sipW-tasA* operon was measured for WT and the 12 Δ*rap-phr* mutants in MSgg under shaking conditions. Only Δ*rapA* showed a significantly reduced relative ON population compared to the WT. Such reduced relative proportion of Δ*rapA* cells in the ON state may be due to more cells committing to activation of the sporulation pathway, and therefore attenuating induction of matrix genes (Bischofs *et al*., 2009). Although only some of the Rap phosphatases have been reported to dephosphorylate Spo0F∼P (Perego, 2013; Omer Bendori *et al*., 2015), we observed that all mutants except Δ*rapA* showed increased matrix gene expression. This indicates that despite the diversity of targets among Rap-Phr systems, and the seeming redundancy of several Rap phosphatases regulating the same master regulator (Perego, 2013), each of the 12 Rap-Phr systems has a regulatory role that affects matrix gene expression under the tested conditions. The involvement of 12 Rap-Phr systems in influencing matrix gene expression suggests that the production of these costly public goods are under complex control, and allows the integration of multiple signals in order to fine-tune the timing of matrix production in response to different conditions (Auchtung *et al*., 2006; Dragoš *et al*., 2018).

Matrix production is well known to be required for formation of architecturally complex biofilms under laboratory conditions (Branda *et al*., 2006; Arnaouteli *et al*., 2021). Moreover, matrix production and localized cell death have been shown to be responsible for the formation of wrinkles during biofilm development (Branda *et al*., 2006; Asally *et al*., 2012; Gallegos-Monterrosa *et al*., 2017). We therefore speculated that the increased matrix gene expression observed for all mutants, except Δ*rapA*, would manifest in these 11 mutants forming more wrinkled colonies as well as more complex, robust pellicles compared to the WT. However, only some of the mutants were affected in biofilm formation. In accordance with increased matrix gene expression, Δ*rapB, J* and *K* mutants formed more wrinkled colonies, and Δ*rapI* and *P* formed complex, though very small colonies compared to the WT. Moreover, Δ*rapC* formed a larger, smoother and more transparent colony. Furthermore, Δ*rapA*, with similar mean expression of the *tapA-sipW-tasA* operon and a smaller relative ON population compared to WT, also formed a more wrinkled colony. Surprisingly, none of the mutants displayed increased pellicle robustness or complexity compared to the WT. In contrast, Δ*rapA, C, I* and *P* formed thinner and/or non-homogenous pellicles.

Next, we were interested in studying how Rap-Phr systems affect biofilm formation of *B. subtilis* in a more ecologically relevant environment, i.e. the plant root. Similarly to biofilm formation *in vitro*, biofilm formation on the plant root by *B. subtilis* depends on matrix gene expression regulated by Spo0A (Beauregard *et al*., 2013; Chen *et al*., 2013). We therefore hypothesized that the increased matrix production observed for most mutants would allow more bacterial cells to attach to and colonize the root. However, only Δ*rapD, J* and *P* showed increased root colonization, while Δ*rapI* was reduced in root colonization. The biofilm and root colonization experiments of the Δ*rap-phr* mutants thus show that the magnitude of matrix gene expression under shaking conditions does not directly correlate with the ability to develop complex biofilms *in vitro* (on agar and at the air-liquid interface) or to colonize the root (Table S1). A lack of positive correlation between matrix gene expression and biofilm formation was observed in a previous study which showed that the magnitude of expression of *epsA-O* and *tasA-sipW-tasA* in *B. subtilis* 168 variants did not directly correlate with formation of wrinkled biofilms (Gallegos-Monterrosa *et al*., 2016). These experiments could thus support that biofilm formation *in vitro* as well as on plant roots is influenced by additional factors than just matrix gene expression. For example surfactin production, which is regulated by ComA – a target of several Rap phosphatases (Perego, 2013) – was shown to influence the colony structure of *B. subtilis* NCIB 3610 on MSgg, though this secondary metabolite was not essential for pellicle formation and root colonization (Thérien *et al*., 2020). However, it has to be noted that matrix gene expression was measured under heavily shaking conditions (220 rpm), while colonies and pellicles were developed under static conditions, and root colonization was performed under mildly shaking conditions (90 rpm). An alternative explanation for the lack of correlation between matrix gene expression under shaking conditions and biofilm formation *in vitro* and on the root could be that the effect of the *rap-phr* deletions on *tasA* expression may vary between these different conditions. Further work is needed to fully explain the discrepancies observed in this study between *tasA* expression and biofilm formation.

Interestingly, several studies have reported a correlation between the ability of strains to form robust biofilms *in vitro* and to colonize the root – both within and among strains (Chen *et al*., 2013; Gallegos-Monterrosa *et al*., 2016). However, the ability of the Δ*rap-phr* mutants to form biofilm *in vitro* did not necessarily reflect the ability to colonize the root (compare Fig. 2 and 3 and summary in Table S1). For example, Δ*rapA* and *C* formed thin and non-homogenous pellicles but were able to colonize the root to similar levels as the WT. Δ*rapD* displayed comparable biofilm *in vitro* to the WT, but was significantly better in root colonization. In addition, Δ*rapJ* formed a highly wrinkled colony, but a pellicle similar to the WT, and was increased in root colonization. Finally, both Δ*rapI* and *P* showed reduced colony size and thin and/or non-homogenous pellicle formation, but while Δ*rapI* showed reduced root colonization, Δ*rapP* was increased in root colonization compared to the WT. These results indicate that the effect of the *rap-phr* deletions on biofilm formation varies between *in vitro* and root conditions. This was similarly shown for a Δ*tagE* mutant (deficient in glycosylating wall teichoic acid) which was affected in root colonization but displayed similar biofilm formation on agar and at the air-liquid interface as the WT (Tzipilevich and Benfey 2021).

In the study by Gallegos-Monterrosa *et al*. (2016), showing that strains forming complex colonies and robust pellicles also efficiently colonize the root, the *B. subtilis* 168 stocks displayed genetic variation in distinct loci (e.g. *epsC* that encodes an enzyme that is directly involved in matrix production), resulting in large differences among the strains in their ability to form biofilm and colonize the root. In contrast, the Δ*rap-phr* mutations studied here might only slightly modulate the regulatory pathways of *B. subtilis*, therefore the ability of the mutants to form biofilm and colonize the root is less altered compared to WT. Nonetheless, the same study also demonstrated that biofilm development is influenced by medium composition (Gallegos-Monterrosa *et al*., 2016). Besides static vs mild agitation and a temperature difference (30 vs 24°C), the media used for testing colony and pellicle formation, and for testing root colonization also slightly differ. First, the MSgg medium used for colony and pellicle formation contains a 10 times higher concentration of glycerol (0.5 %) compared to the MSNg medium used for plant root colonization (0.05 %). During plant root colonization, the bacteria thus depend on plant polysaccharides and root exudates as carbon source. In addition, *in vitro* biofilm development depends on the availability of iron and manganese (Kolodkin-Gal *et al*., 2013; Shemesh and Chai, 2013; Mhatre *et al*., 2016), while during plant root colonization in MSNg, biofilm formation is induced by plant polysaccharides (Beauregard *et al*., 2013). The presence of plant polysaccharides may therefore allow Δ*rap-phr* mutants that form weak pellicle biofilms *in vitro* to efficiently colonize the root *(ΔrapA, C* and *P*, Fig. 2 and 3). Thereby, the pathways regulating matrix gene expression and other factors important for biofilm formation may be differently regulated under distinct conditions.

Finally, the disparate results obtained in this study may be understood in the light of the full set of 12 Rap-Phr systems with redundant functions: the influence of a single *rap-phr* deletion might be masked by the function of another redundant Rap-Phr system. Furthermore, if such potential redundancy varies between the different conditions employed in this study (e.g. if the *rap*-*phr* genes are differentially expressed under the distinct conditions tested), this could (partly) explain the observed discrepancy, e.g. between biofilm formation *in vitro* and on the root.

To conclude, we here show that all 12 Rap-Phr systems have an impact on matrix gene expression in liquid culture. Thereby, the diversity in Rap-Phr systems in *B. subtilis* 3610 could function to integrate multiple signals in order to fine-tune the timing and level of responses to new ecological niches, such as those it will encounter in soil. Furthermore, we show that the ability to form biofilm *in vitro* not necessarily reflects the ability to colonize the root under the tested conditions. These findings thus support that the pathways involved in matrix gene expression and other components important for biofilm establishment could be differently influenced under distinct conditions.

## Experimental procedures

### Bacterial strains and cultivation methods

Strains used in this study are listed in Table 1. The *B. subtilis* DK1042 strain (a natural competent derivative of the undomesticated NCBI 3610)(Konkol *et al*., 2013) was used as WT. The 12 single Δ*rap-phr* mutants were previously created and contains an antibiotic resistance cassette in place of the *rap-phr* gene pair (except for the markerless *ΔrapB* mutant) (Gallegos-Monterrosa *et al*., 2020). For flow cytometry, the Δ*rap-phr* mutants were transformed with the plasmid phy_mKATE2 harboring the *mKATE* gene under the control of the hyper-spank promoter (which is constitutive due to the absence of *lacI*) and a chloramphenicol (Chl) resistance gene within the flanking regions of the *amyE* gene (van Gestel *et al*., 2014). Transformants were identified by selecting for Chl resistance, and double cross overs were verified by the loss of amylase activity. The resulting mKATE-labelled Δ*rap-phr* mutants were transformed with genomic DNA from *B. subtilis* TB373, which harbors the P_*tapA*_*-gfp* reporter construct with a kanamycin (Km) resistance gene integrated at the *sacI* locus. Successful transformants with the reporter construct inserted into the *sacI* locus were identified by selecting for Km resistance. The resulting reporter strains were verified for reporter activity under the fluorescence stereomicroscope. Importantly, due to the presence of a Km resistance cassette in place of the corresponding *rap-phr* gene pair in Δ*rapA, C, D, I, J* and *K*, markerless versions of those *rap-phr* mutants were used to obtain the reporter strains for the *tapA-sipW-tasA* operon for the flow cytometry analysis. For all experiments, strains were grown overnight in Lysogeny broth (LB; Lennox, Carl Roth; 10 g×l^-1^ tryptone, 5 g×l^-1^ yeast extract and 5 g×l^-1^ NaCl) at 37°C while shaking (220 rpm). For transformation and stock preparation, antibiotics were used at the following working concentrations: Km: 5 µg×ml^-1^ and Chl: 5µg×ml^-1^. For analyzing biofilm formation and matrix gene expression, strains were grown in MSgg (5 mmol×l^-1^ potassium phosphate [pH 7], 0.1 mol×l^-1^ 3-(N-morpholino)propanesulfonic acid (MOPS) [pH 7], 2 mmol×l^-1^ MgCl_2_, 700 μmol×l^-1^ CaCl_2_, 100 μ×l^-1^ MnCl_2_, 50 μmol×l^-1^ FeCl_3_, 1 μmol×l^-1^ ZnCl_2_, 2 μmol×l^-1^ thiamine, 0.5% glycerol, and 0.5% K-glutamate). For root colonization assay, strains were grown in MSNg (5 mmol×l^-1^ potassium phosphate buffer [pH 7], 0.1 mol×l^- 1^ MOPS [pH 7], 2 mmol×l^-1^ MgCl_2_, 50 μmol×l^-1^ MnCl_2_, 1µmol×l^-1^ ZnCl_2_, 2 μmol×l^-1^ thiamine, 700 μmol×l^-1^ CaCl_2_, 0.2 % NH_4_Cl_2_, and 0.05 % glycerol).

**Table 1:**
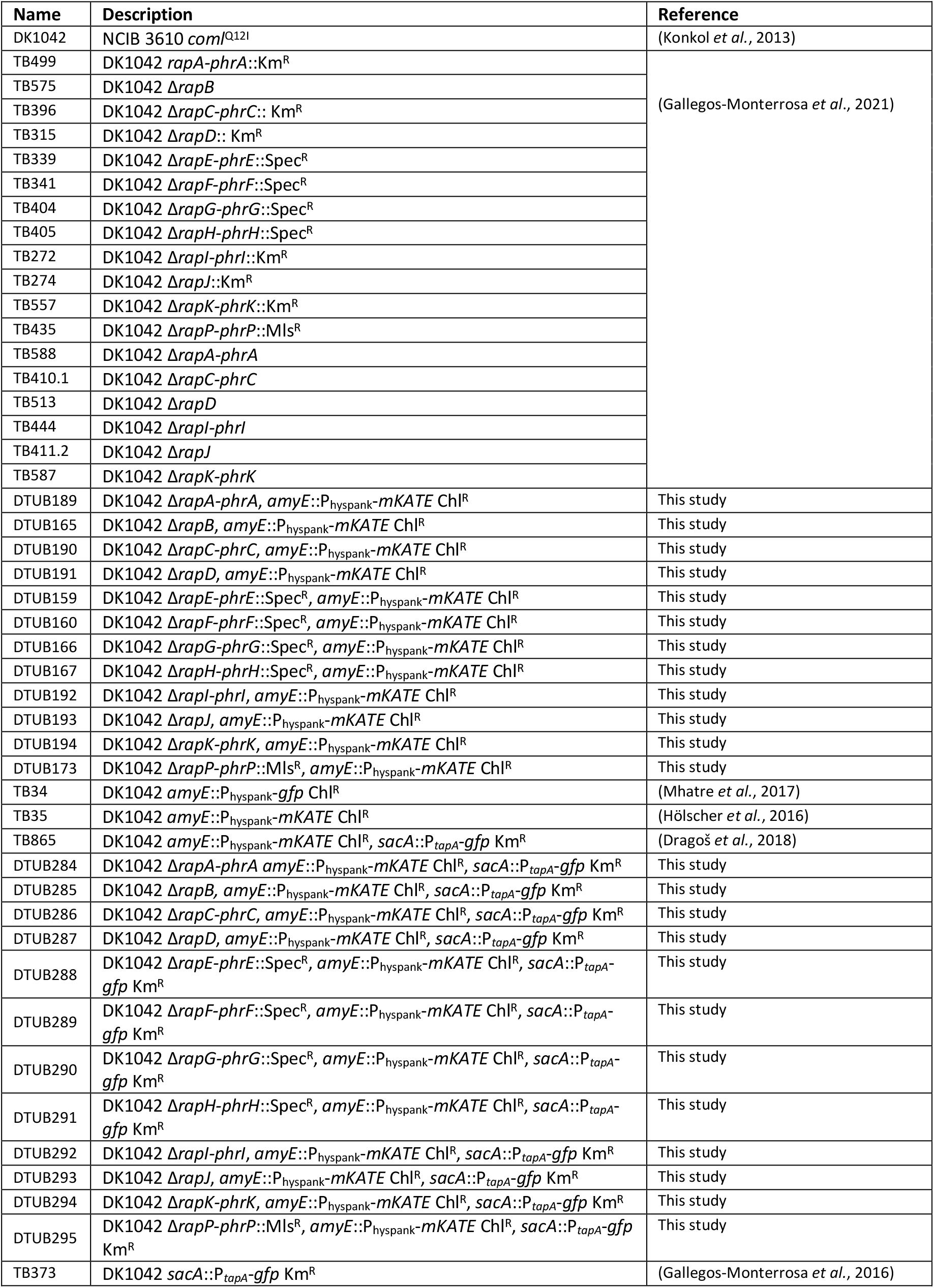
Strains used in this study.

### Microscopy imaging

All images were acquired with an Axio Zoom V16 stereomicroscope (Carl Zeiss, Germany) equipped with a Zeiss CL 9000 LED light source and an AxioCam MRm monochrome camera (Carl Zeiss, Germany).

### Biofilm formation assay

For biofilm formation on a solid surface, 7 µL overnight cultures were spotted on MSgg supplemented with 1.5 % agar. For pellicle biofilm formation at the air-liquid interface, 15 µl inoculum of overnight cultures adjusted to OD_600_ of 5 was added to 1.5 ml MSgg medium in 24-well plates, giving a starting OD_600_ of 0.05. Plates were incubated under static conditions at 30°C for 48 h, thereafter images of the arisen colonies and pellicles were obtained using the stereomicroscope.

### Root colonization assay

*A. thaliana* Col-0 plants were used as a host for *B. subtilis* root colonization. *A. thaliana* seeds were sterilized in 2% (v/v) sodium hypochlorite (NaOCl) for 10 minutes with an orbital shaker. Following this, NaOCl was removed and the seeds were washed five times in sterile water. Sterilized seeds were placed in MS agar plates (Murashige and Skoog basal salts, Sigma) (2.2 g×l^-1^) with approximately 20 seeds per petri dish. The petri dishes were wrapped in parafilm and left for stratification at 4°C for 3 days in order to break seed dormancy and were then moved to the plant chamber (cycles of 16 h light at 24°C and 8 h dark at 20°C). After five to seven days, seedlings of 0.5-1.2 cm in size were placed in 48-well plates containing 270 µl MSNg medium per well. To each well, 30 µl of overnight culture adjusted to OD_600_ of 0.2 was added resulting in a final starting OD_600_ of 0.02. The plates were sealed with parafilm and incubated in the plant chamber while shaking at 90 rpm for 16 h. Seedlings were then washed in MSNg to remove non-attached cells from the root. The washed seedlings were placed in Eppendorf tubes containing 1ml of NaCl (0.9 %) and subjected to standard sonication protocol to disperse the biofilm (Dragoš *et al*., 2018). The resulting bacterial cell suspension was diluted and plated on LB agar plates for CFU counting. To acquire the CFU per mm root the obtained CFU was divided by the length of the corresponding root.

### Growth profiling

To follow the growth of WT and mutants, two overnight cultures of each strain were independently inoculated into a 96-well plate containing MSgg or LB broth, at a starting OD_600_ of 0.05. Growth was monitored in a plate reader (Infinite F200 PRO, TECAN and BioTek Synergy HTX Multi-Mode Microplate Reader) every 10 min for 48 h at 30°C under linear shaking conditions.

### Matrix gene expression assays by flow cytometry

To measure expression from the matrix operon *tapA-sipW-tasA*, mKate-labelled WT and Δ*rap-phr* mutants harboring the reporter construct P_*tapA*_*-gfp* were incubated in 10 ml MSgg in 100 ml bottles with a starting OD_600_ of 0.02. Lids were loosely on, so oxygen would not be depleted. Bottles were incubated at 37°C at 220 rpm for 6 h. After incubation, 1 ml of each sample was transferred to a 2 ml Eppendorf tube and samples were run on the flow cytometer (MACSQuant® VYB, Miltenyibiotec). mKate-positive cells were detected by yellow laser (561 nm) and filter Y3 (661/20 nm). Green fluorescent cells, representing the cells expressing the *tapA-sipW-tasA* operon, were detected by the blue laser (488 nm) and filter B1 (525/50 nm). Strain TB35 with constitutive mKate expression was used as a negative control for GFP expression, while strain TB34 with constitutive GFP expression was used as a positive control for GFP expression. In addition, TB34 and a medium control were used to identify the red background fluorescence noise due to autofluorescence, and cells above this background fluorescence were identified as the mKate positive cells representing *B. subtilis* cells producing the mKate protein. For each WT and mutant sample, single events were identified on the SSC-H vs SSC-A plot and gated into the mKate-A vs SSC-A plot, where mKate positive cells were identified. These gated cells were exported and read into Excel where the green fluorescence (GFP-A) values were used for analysis. To get rid of negative values, 300 AU was added to all events in the samples. To obtain the distribution of GFP expression, data obtained from each replicate was subjected to binning with identical bin size (of 50). Events with GFP expression from 0 to 10.000 were included as the majority of events were within this interval (>98 %). Next, the number of events in each bin was divided by the total number of events in the given replicate, resulting in the normalized frequency. To obtain the mean distribution, a mean frequency for each bin was obtained by averaging the individual frequencies within this bin across the replicates, resulting in the mean distribution of single-cell-level expression. For statistics, the relative OFF (GFP values between 0 and 500) and ON (GFP values between 500-10.000) populations were calculated for each replicate, as was the mean fluorescence of the ON population.

### Statistical analysis

Statistical analyses were performed in R Studio. For each root colonization assay, a one-way analysis of variance (ANOVA) was performed on the log10-transformed data. P-values were adjusted using the Benjamini & Hochberg procedure. When adjusted P-values were significant (P<0.05), a Dunnett’s Multiple Comparison test was performed. When data failed to meet parametric assumptions (normality and equal variance), a non-parametric analysis (Kruskal-Wallis test) was used. If adjusted P-values were significant (P<0.05), treatment means were compared via a Dunn test. To test for significant differences in relative OFF/ON population and mean fluorescence of ON population between mutants and WT, an ANOVA followed by Dunnett’s Multiple Comparison test was performed. Significance level for all tests were set at 5 %.

## Acknowledgements

We thank Mikael Lenz Strube for his suggestions on statistics. The work was supported by a DTU Bioengineering start-up fund to ÁTK. Funding from Novo Nordisk Foundation (grant NNFOC0055625) for the infrastructure “Imaging microbial language in biocontrol (IMLiB)” is acknowledged.

## Authors’ contributions

M.N.C. and Á.T.K. conceived the project; M.N.C., R.M.R.M., N.K.K. performed experiments; R.G.-M. created mutant strains; M.N.C. analyzed the data; M.N.C. and Á.T.K. wrote the manuscript with feedback from all authors.

