## Supplementary material for "Deletion of Rap-Phr systems in *Bacillus subtilis* influences *in vitro* biofilm formation and plant root colonization": Fig S1 to S2 and Table S1

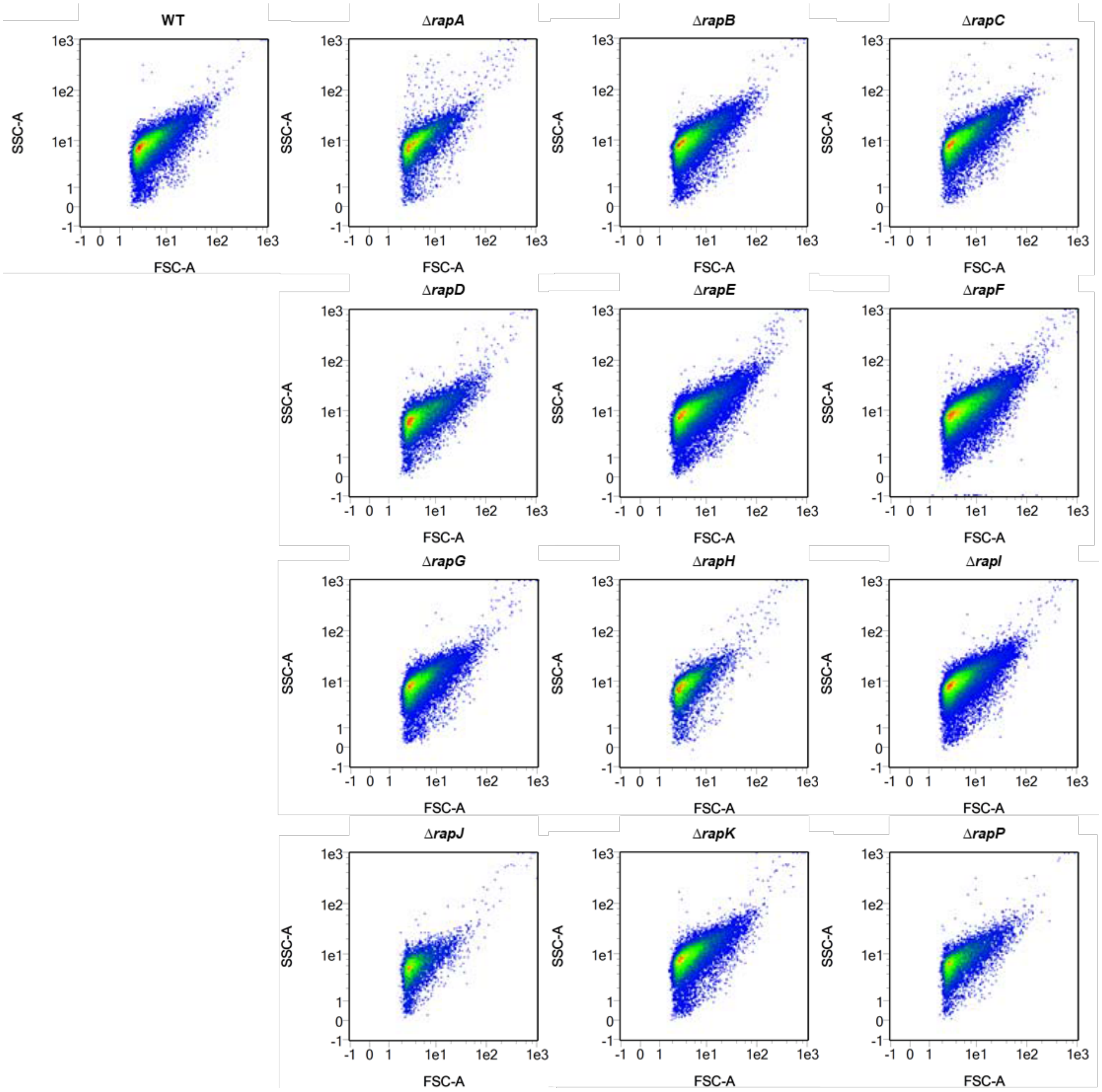


**Fig. S1: Single Δ*rap-phr* mutants are not majorly affected in cell granularity or size.** Flow cytometry analysis showing the side scatter (SSC-A) vs forward scatter (FSC-A) plots of ungated WT and *Δrap-phr* mutant cells harboring the $P_{tapA}$-*gfp* construct.

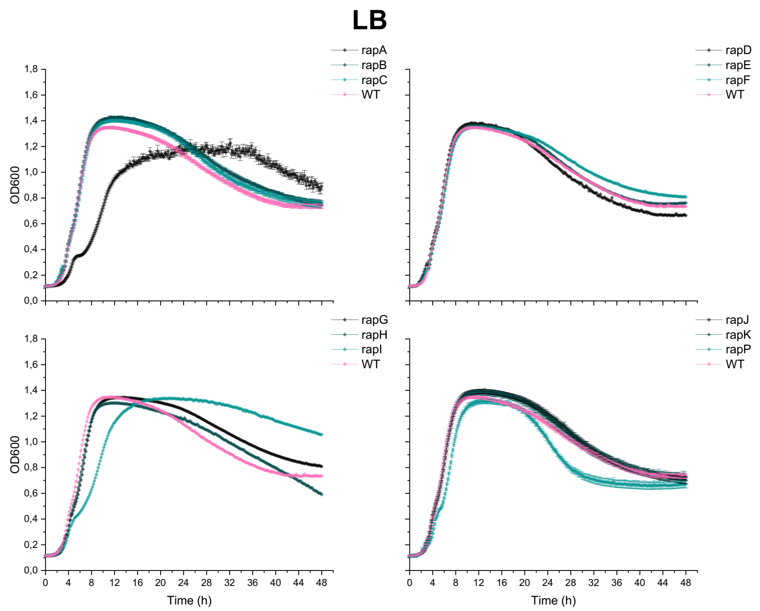


**Fig. S2: Growth of *B. subtilis* WT and single Δ*rap-phr* mutants in LB medium.** WT and mutants were inoculated into 96-well plates with a starting $OD_{600}$ of 0.05. $OD_{590}$ was measured every 10 min for 48 h at 30°C, and each time point represents the mean of six technical replicates from two overnight cultures (n=6). Error bars represent standard error (SE).

**Table S1: Overview of phenotypes of the 12 single *Δrap-phr* mutants compared to WT.** Up arrows indicate increased while down arrows indicate decreased features compared to WT. For colony and pellicle formation, the direction of the arrow is related to wrinkles and complexity.

| Mutant | Known target of respective Rap proteins | Matrix expr. | Colony | Pellicle | Root | Growth |
|---|---|---|---|---|---|---|
| *rapA-phrA* | Spo0F | ↓ | ↑ | ↓ | - | ↓ |
| *rapB* | Spo0F | ↑ | ↑ | - | - | - |
| *rapC-phrC* | ComA | ↑ | ↓ | ↓ | - | - |
| *rapD* | ComA | ↑ | - | - | ↑ | - |
| *rapE-phrE* | Spo0F | ↑ | - | - | - | - |
| *rapF-phrF* | ComA | ↑ | - | - | - | - |
| *rapG-phrG* | DegU | ↑ | - | - | - | - |
| *rapH-phrH* | Spo0F, ComA | ↑ | - | - | - | - |
| *rapI-phrI* | Spo0F, regulation of mobile genetic elements | ↑ | ↑ (but small) | ↓ | ↓ | ↓ |
| *rapJ* | Spo0F | ↑ | ↑ | - | ↑ | - |
| *rapK-phrK* | ComA | ↑ | ↑ | - | - | - |
| *rapP-phrP* | Spo0F, ComA | ↑ | ↑ (but small) | ↓ | ↑ | ↑ |
